# Vancomycin Relieves Mycophenolate Mofetil-Induced Gastrointestinal Toxicity by Eliminating Gut Bacterial β-Glucuronidase Activity

**DOI:** 10.1101/561274

**Authors:** Michael R. Taylor, Kyle L. Flannigan, Hannah Rahim, Amina Mohamud, Ian A. Lewis, Simon A. Hirota, Steven C. Greenway

## Abstract

Mycophenolate mofetil (MMF) is commonly prescribed after transplantation and has proven advantages over other immunosuppressive drugs but gastrointestinal (GI) side effects frequently limit its use. The pathways involved in the metabolism of the prodrug MMF and the bioactive derivative mycophenolic acid (MPA) are well characterized but the mechanism responsible for toxicity is unknown. Here we extend our previous observation that an intact gut microbiome is required for MMF-induced toxicity and demonstrate that gut bacterial metabolism is responsible for the GI inflammation and weight loss associated with MMF exposure. In mice consuming MMF, the introduction of vancomycin alone was sufficient to prevent or reverse MMF-induced weight loss and colonic inflammation. MMF induced the expansion of bacteria expressing β-glucuronidase (GUS) in the cecum and proximal colon. GUS activity, which is responsible for the catabolism of glucuronidated MPA (MPAG) to free MPA, was increased in the presence of MMF and eliminated by vancomycin. Vancomycin eliminated multiple *Bacteroides spp*. that flourished in the presence of MMF and prevented the breakdown of MPAG without negatively affecting serum MPA levels. Human data suggests that increased stool GUS activity can be associated with MMF-related toxicity. Our work provides a mechanism for the GI toxicity associated with MMF and a future direction for the development of therapeutics.

## Introduction

Suppression of the immune system is essential after hematopoietic stem cell or solid organ transplantation and is commonly used for the treatment of multiple autoimmune diseases. One of the most widely used immunosuppressive drugs is mycophenolate mofetil (MMF) (*1-4*). MMF has important advantages over other drugs (*5-8*) but, while effective in enhancing survival, frequent gastrointestinal (GI) side effects can lead to dose reduction or discontinuation, actions linked to increased risk of rejection and patient death after solid organ transplantation (*9-14*). The xenobiotic metabolism of MMF within humans has been well characterized (Fig. 1) (*15*). MMF is hydrolyzed to active mycophenolic acid (MPA) in the stomach and small intestine and then inactivated by hepatic uridine diphosphate glucuronosyltransferase (UGT) enzymes to create glucuronidated MPA (MPAG) and a small amount of acyl-MPAG (AcMPAG). Enteric-coated MMF and intravenous MMF are also rapidly converted to MPA and subsequently glucuronidated in the liver. MPAG and AcMPAG are excreted primarily in the urine but ∼10% is transported by the ATP-binding cassette sub-family C member 2 (ABCC2) protein into the GI tract where bacteria expressing the enzyme β-glucuronidase (GUS) remove the glucuronic acid (GA) moiety to produce free MPA and GA (*16*). The resulting GA is available as a carbon source for bacterial metabolism and the MPA undergoes enterohepatic recirculation.

**Fig. 1.**
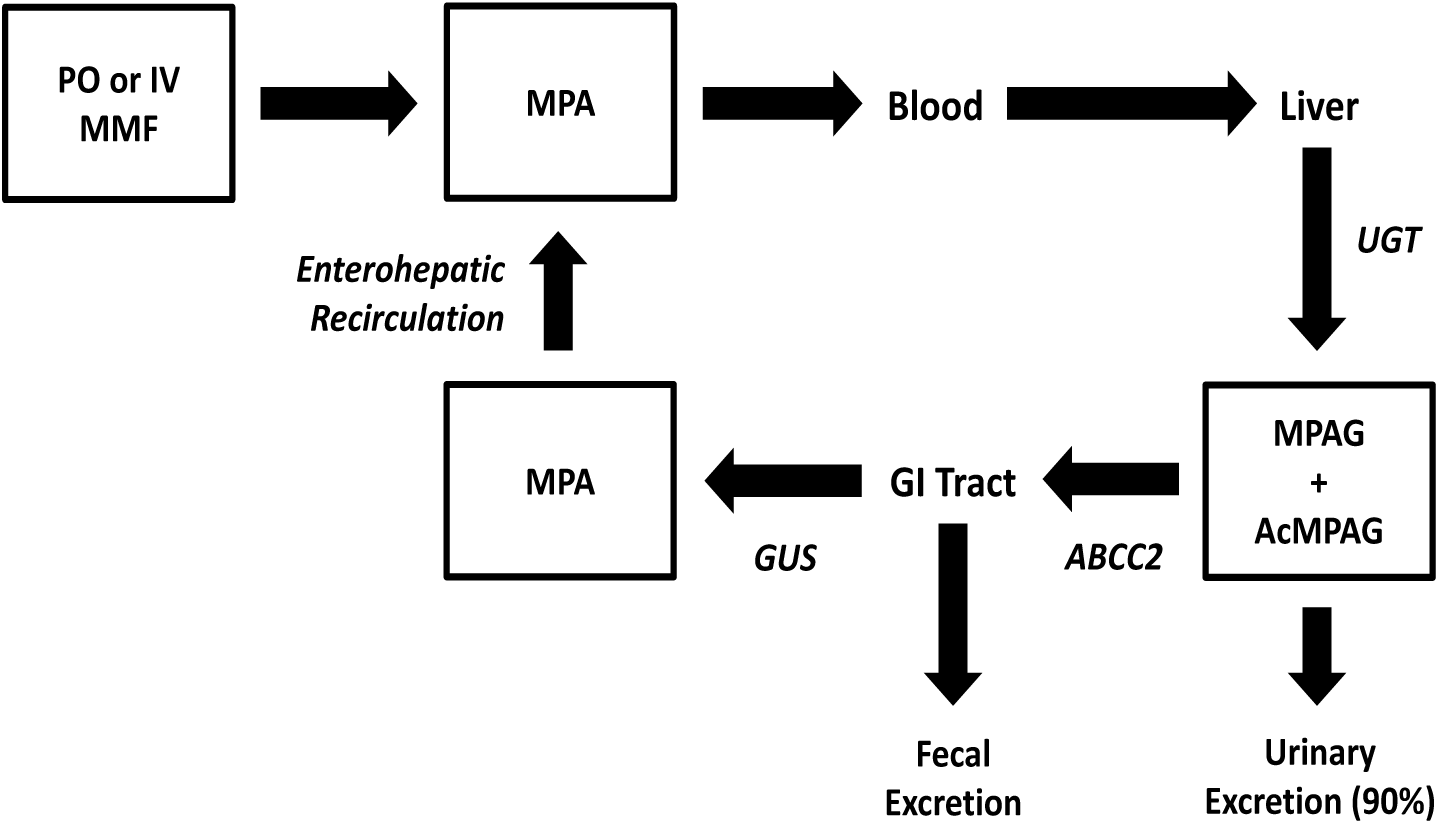
MMF metabolism. MMF consumed either orally (PO) or intravenously (IV) is converted to active MPA in the stomach and circulation and then glucuronidated by hepatic UGT enzymes which create inactive MPAG and AcMPAG. MPAG and AcMPAG are mostly excreted in the urine but ∼10% is transported by ABCC2 into the GI tract where bacterial GUS enzymes remove the GA moiety to produce free GA and MPA which undergoes enterohepatic recirculation.

The gut microbiota, referring to all of the microorganisms in this niche, and its associated metagenome (microbiome) influences diverse aspects of human physiology and changes in its composition impacts human health (*17, 18*). The mechanism(s) contributing to MMF-related GI toxicity have yet to be fully characterized but we have shown that hematochezia and colonic inflammation in a pediatric heart transplant patient taking MMF was associated with a shift in the composition of the intestinal microbiota and that mice consuming MMF develop weight loss, colonic inflammation and changes in their gut microbiota (*19, 20*). In our patient, toxicity only resolved after the discontinuation of MMF but, in our mouse model, MMF-induced GI toxicity was reversed or prevented by the use of broad-spectrum antibiotics and was absent in germ-free animals. These observations suggest that MMF toxicity requires an intact gut microbiota. In the current study, we sought to further investigate the role of gut bacteria in MMF-induced GI toxicity using our mouse model. We hypothesized that MMF toxicity would be associated with alterations in the composition and function of the microbiome. We identified the ability of a single antibiotic, vancomycin, to both prevent and reverse the pathophysiological effects induced by MMF consumption and we characterized the impact of vancomycin on the gut microbiome. We then explored the activity of gut bacterial GUS enzymes in the presence of MMF and vancomycin and examined the impact of vancomycin on MPAG catabolism. Our data suggest that the introduction of MMF alters the composition of the gut microbiota by selecting for GUS-expressing bacteria. Increased GUS expression and activity leads to increased concentrations of MPA in the colon which is associated with colonic inflammation and weight loss. Human data suggests that increased stool GUS activity can be associated with MMF-related toxicity. Vancomycin eliminated the GUS-expressing bacteria and prevented MMF-induced weight loss and colonic inflammation without reducing therapeutic levels of MPA in the blood. This work demonstrates a mechanism for MMF-induced GI toxicity and suggests a potential clinical intervention. The interaction between MPA and the gut microbiome provides further insight into the significant problem of drug intolerance and adverse drug reactions driven by microbial metabolism and may also represent a paradigm for how the metabolism of xenobiotics can create local GI pathology.

## Results

### Vancomycin is sufficient to prevent and reverse MMF-induced weight loss

Using an established mouse model of MMF exposure, we found that vancomycin alone, when administered concurrently with MMF, prevented the weight loss caused by MMF treatment (Fig. S1). The weight of vancomycin-treated animals did not differ significantly from control, untreated animals after 8 days of MMF exposure (103.2 ± 0.68% vs. 98.84 ± 0.82%, p = 0.41). Vancomycin was also able to reverse MMF-induced weight loss even with continued exposure to MMF. As seen previously, MMF caused significant weight loss with animals weighing 75.02 ± 1.5% of their initial weight after 8 days, but, in the subsequent 8 days after the introduction of vancomycin, animal weights improved significantly with recovery back to 93.82% ± 1.2% of their initial weight (Fig. 2A). Although animals exposed to MMF and vancomycin were still significantly smaller than control, untreated animals (105.7 ± 1.5%, p = 0.003) they were significantly heavier than animals that continued to be exposed to MMF (69.26 ± 0.93%, p = 0.014). Consistent with the improvement in body weight, vancomycin also improved colon length and restored weight of the cecum, liver, and spleen. However, hematocrit remained significantly reduced (Fig. S2). Whole colonic tissue homogenates were used to assess changes in the cytokine and chemokine profile of treated mice. Mice consuming both MMF and vancomycin demonstrated significant decreases (p < 0.05) in the levels of KC, G-CSF, IL-6, MIP-2, LIF, TNFα and M-CSF and significant increases (p < 0.05) in IL-2, RANTES, MIG and IP-10 (Fig. 2B and Fig. S3). In addition to vancomycin, the effects of individual neomycin, ampicillin and metronidazole were tested for their ability to reverse MMF-induced weight loss (Fig. S1). Metronidazole treatment actually exacerbated the MMF-induced weight and ampicillin and neomycin were less reproducibly effective than vancomycin in preventing MMF-induced weight loss.

**Fig. 2.**
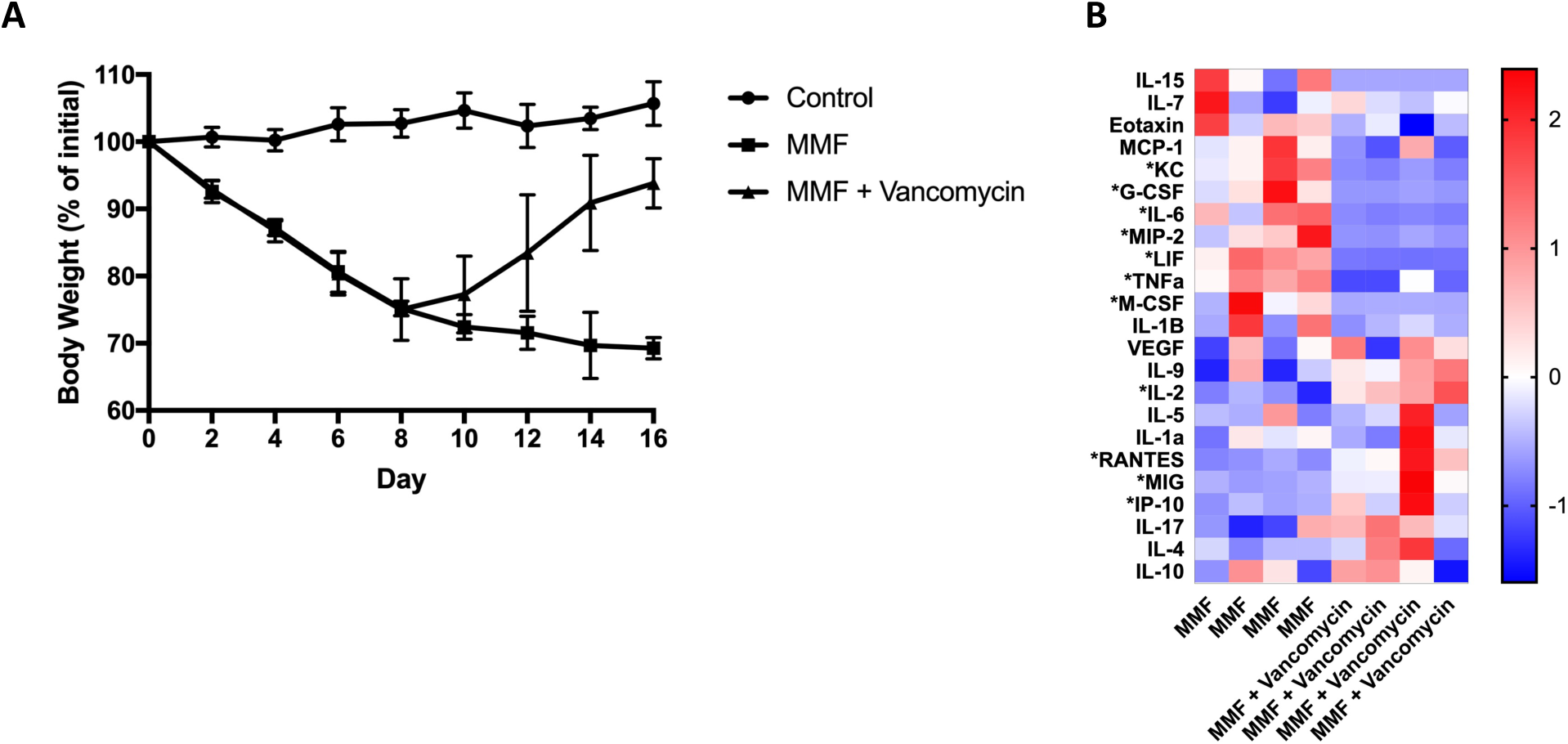
Effect of vancomycin on mouse body weight and colonic inflammation. (**A**) Vancomycin was sufficient to reverse MMF-induced weight loss even in the continued presence of MMF. Data represent two separate experiments with 4-9 animals for each group and are expressed as mean and standard deviation for each time point. (**B**) Vancomycin induced significant changes (*, p <0.1) in multiple colonic cytokines and chemokines. Each column represents an individual animal.

### Vancomycin abolishes GUS-producing gut bacteria

Using 16S rRNA amplicon sequencing, we found vancomycin to have broad effects on the composition of the gut microbiota. After two days of vancomycin treatment (day 10), the gut microbiome of mice treated with both MMF and vancomycin was significantly (p < 0.05) distinct from the MMF-only group with regards to β-diversity as measured using a non-metric multidimensional scaling (NMDS) plot of Bray-Curtis dissimilarity and continued to separate from the MMF group over time (Fig. S4). After treatment with vancomycin, the α-diversity of the gut microbiome decreased significantly as measured by observed species (277 vs. 549, p = 2.8 × 10^-4^), Shannon diversity index (1.42 vs. 3.83, p < 1.0 × 10^-7^) and the Simpson’s diversity index (0.59 vs. 0.92, p = 8.0 × 10^-4^) after only 2 days of antibiotic exposure. The Shannon and Simpson’s diversity indices both continued to be significantly lower in the MMF and vancomycin-treated group for the following 6 days (Fig. S4).

At the class level, dominant bacteria included Clostridia, Bacteroidia and Bacilli during the initial eight days of MMF treatment (Fig. 3A and Fig. S5A). Continued MMF exposure led to an expansion of Gammaproteobacteria, Erysipelotrichia and, to a lesser extent, Deltaproteobacteria. After the addition of vancomycin on day 8, the Bacteroidia, Clostridia, and Erysipelotrichia taxa were severely diminished, whilst the Bacilli and Gammaproteobacteria flourished. By Day 16, after 8 days of combined vancomycin and MMF exposure, the bacterial community was composed almost entirely of class Bacilli with only a small proportion of Gammaproteobacteria remaining as the next predominant class. The classes Bacilli and Betaproteobacteria were significantly increased (p < 0.05) after the addition of vancomycin (Fig. S5A). Considering the differentially depleted classes (Bacteroidia, Clostridia and Erysipelotrichia present in the MMF-treated animals but eliminated by vancomycin), the genera that were both significantly differentially affected by vancomycin and in relatively high abundance in animals after 8 days of MMF treatment were Bacteroides, Lachnospiraceae NK4A136, Roseburia, and Turicibacter (Fig. S5B). We generated species-level resolution for Bacteroides including *B. vulgatus*, *B. fragilis*, *B. caccae*, *B. uniformis*, *B. ovatus* and *B. nordii*, all of which were detectable in the presence of MMF but rapidly eliminated by vancomycin (Fig. 3B).

**Fig. 3.**
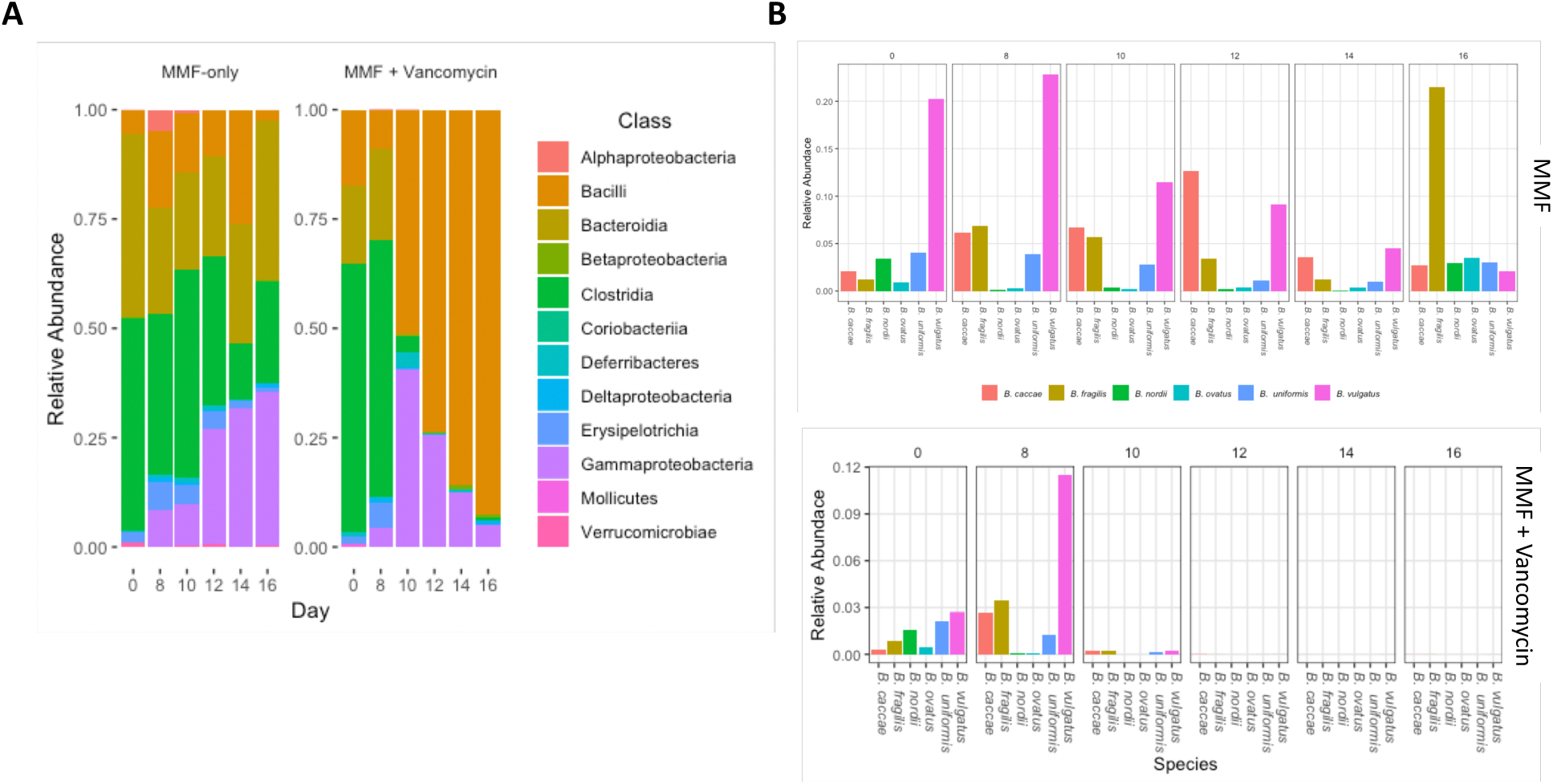
Vancomycin-induced changes in the gut microbiome. Amplicon sequencing of DNA extracted from mouse fecal pellets was used to determine bacterial composition. (**A**) Stacked bar plots showing relative abundance of bacterial classes in the presence of MMF with and without vancomycin. Vancomycin decreased overall diversity with a gradual predominance of Bacilli. (**B**) *Bacteroides* spp. were abundant in the presence of MMF but were rapidly eliminated by the introduction of vancomycin on day 8.

### GUS gene expression and actvity are increased by MMF and abolished by vancomycin

We examined the proportional abundance of the bacterial GUS gene using PICRUSt. We found the predicted proportional abundance of GUS enzyme orthologs (KEGG Orthology K01195, KEGG Enzyme EC 3.2.1.31) to be comparable between groups before commencing MMF treatment. After 4 days of MMF treatment, the MMF-only group had a significant increase in the predicted proportional abundance of K01195 (p = 0.006) but, after 8 days, the abundance in the MMF-only group had decreased slightly (Fig. 4A). Functionally, the activity of GUS enzymes, assessed in fecal pellets of untreated and MMF-treated mice, followed the same pattern, with the treated mice exhibiting a significant increase in enzyme activity after 4 days of exposure (Fig. 4B). During the initial 8 days of MMF-only treatment, prior to the initiation of vancomycin, there were no differences between the two groups with regards to gene expression (p = 0.33) (Fig. 4C). The introduction of vancomycin after 8 days of MMF treatment reduced the predicted proportional abundance of K01195 after only 2 days (day 10). After 8 days of vancomycin treatment (day 16), the abundance of K01195 was reduced to nearly undetectable levels, in comparison to pre-antibiotic levels (p = 1.2 × 10^-7^), despite continuation of the MMF exposure. Concordant with reduced gene expression, GUS activity in fecal pellets was abolished by treatment with vancomycin (Fig. 4D).

**Fig. 4.**
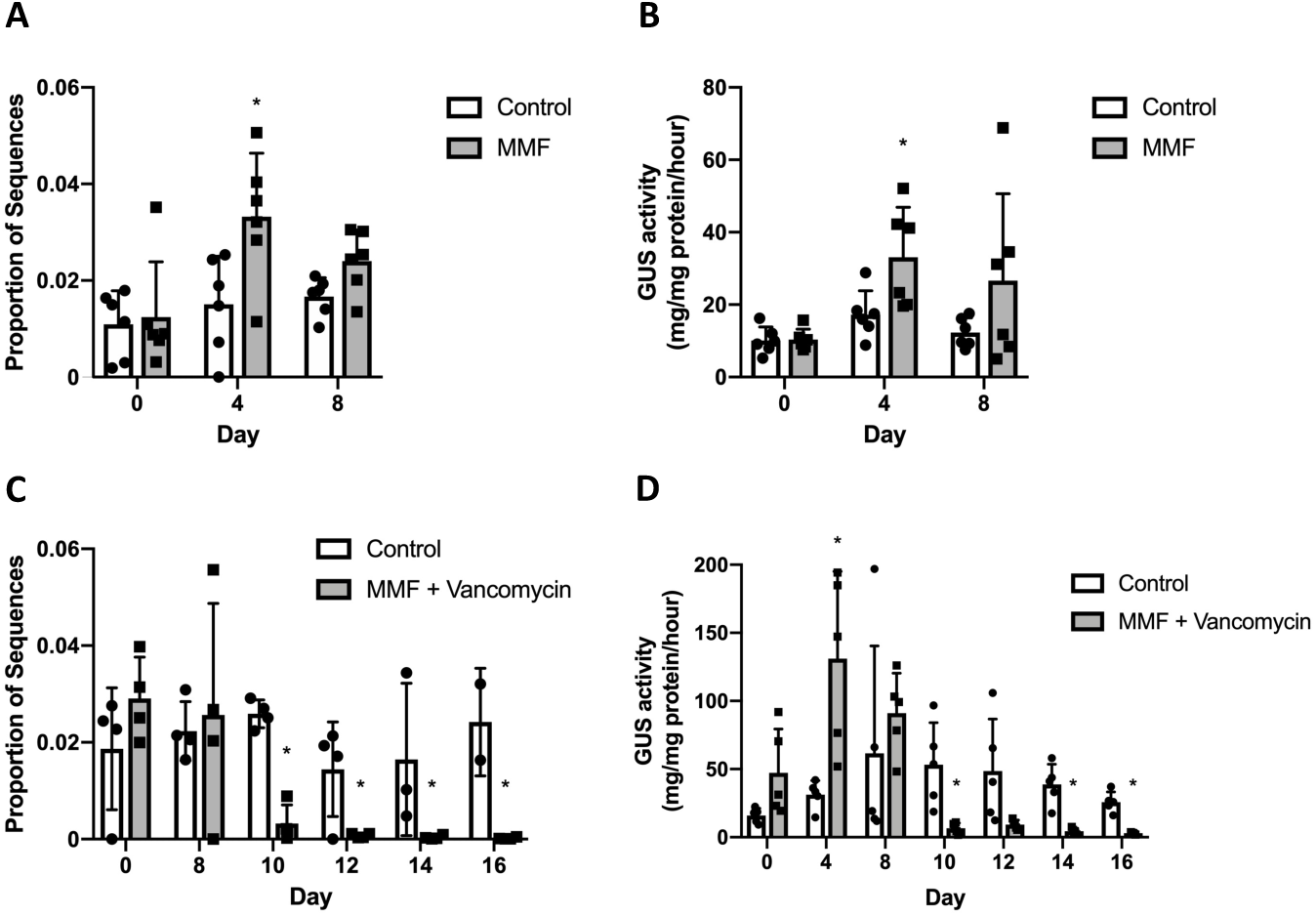
MMF-upregulated GUS expression and activity is prevented by vancomycin. PICRUSt and an enzyme activity assay were used to assess gene expression and GUS activity in mouse fecal pellets. (**A**) The proportional abundance of KEGG ortholog K01195 increased significantly (*p = 0.006) in the presence of MMF after 4 days. (**B**) GUS activity in fecal pellets collected from mice consuming MMF also increased significantly (p = 0.03) after 4 days. (**C**) The introduction of vancomycin on day 8 rapidly eliminated the expression of K01195. (**D**) Vancomycin, introduced after day 8, abolished GUS activity in fecal pellets (*p<0.01). All data are plotted as means and standard deviations with data points for individual samples represented.

### In vivo imaging of intestinal GUS activity

The effect of MMF and vancomycin on intestinal GUS activity in the intact animal was assessed using the fluorescent substrate FDGlcU. Mice gavaged with FDGlcU and exposed to MMF demonstrated an upregulation of GUS activity in the proximal colon after 8 days of treatment but not in the cecum as compared to controls. Vancomycin significantly reduced GUS activity in both the cecum (day 4, p = 0.05; day 8, p = 0.008) and proximal colon (day 8, p = 0.03) and also caused a qualitative enlargement of the cecum (Fig. 5).

**Fig. 5.**
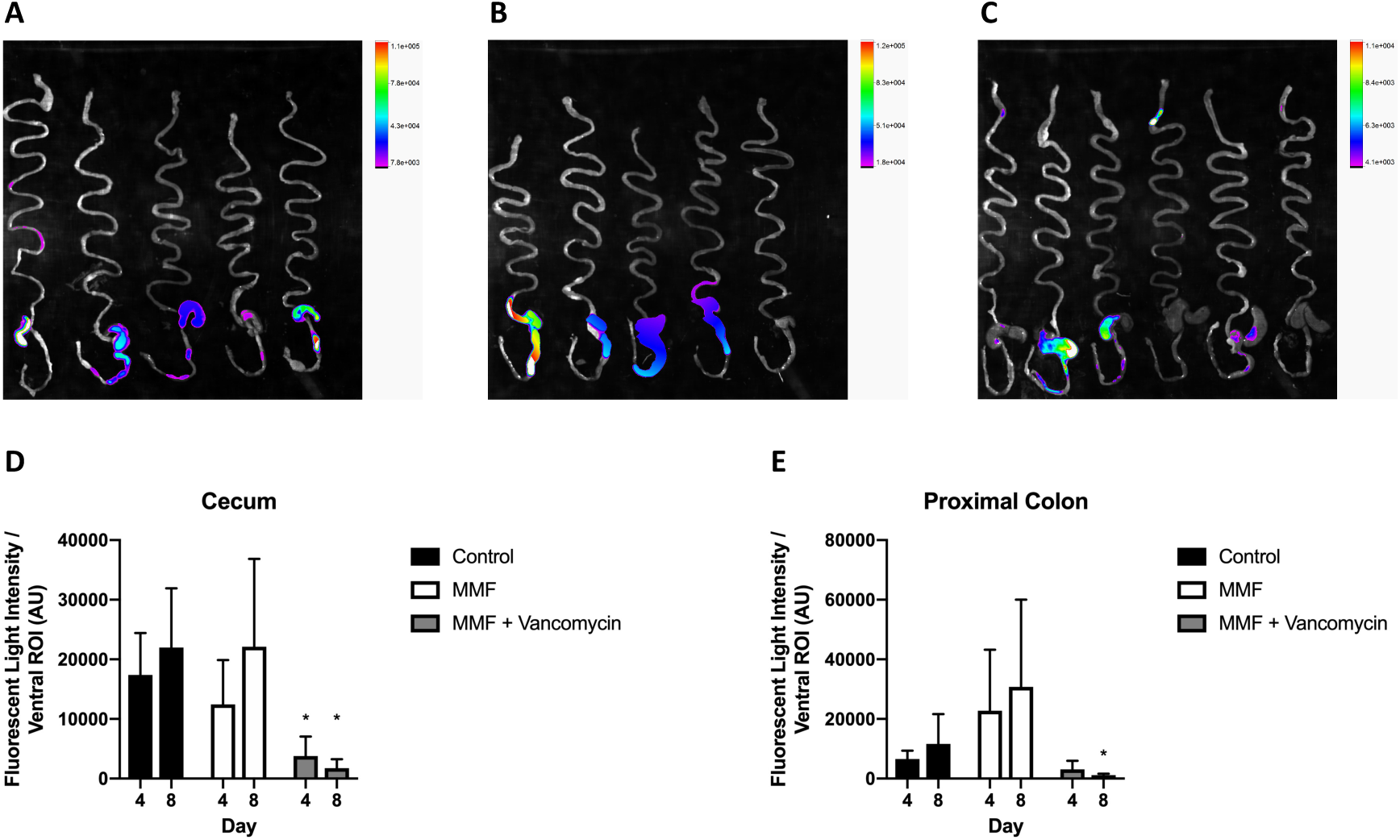
Intestinal GUS activity is decreased by vancomycin. Mice were treated with MMF or MMF and vancomycin for 8 days and then GUS activity was imaged using the fluorescent substrate FDGlcU. (**A**) *Ex vivo* imaging of the GI tract from control mice. (**B**) *Ex vivo* imaging of the mouse GI tract after animals consumed MMF for 8 days. (**C**) *Ex vivo* imaging of the GI tract of mice exposed to MMF and vancomycin for 8 days. (**D**) Quantification of GUS activity in the cecum did not demonstrate a significant increase with MMF exposure after 4 or 8 days of exposure but activity decreased significantly after 4 and 8 days of vancomycin treatment. (**E**) Quantification of GUS activity in the proximal colon did not demonstrate a significant increase with MMF but activity decreased significantly after 8 days of vancomycin treatment compared to animals receiving MMF only. *p < 0.05.

### Vancomycin limits MPAG catabolism

The catabolism of MPAG/AcMPAG (undifferentiated in our fecal assay) to MPA and GA was found to be significantly reduced by the addition of vancomycin in MMF-treated mice, as measured in fecal pellets using UHPLC-MS (Fig. 6A). During the MMF-only exposure (Days 0 to 8), we observed relatively high average concentrations of MPA in both groups (35.2 ± 4.13 μM and 33.73 ± 4.50 μM) and relatively low average concentrations of MPAG (1.85 ± 0.48 μM and 1.53 ± 0.47 μM). After the addition of vancomycin on day 8, there was a significant increase in the average concentration of MPAG (16.01 ± 0.92 μM, p = 0.01) and a nearly 8-fold decrease in the average MPA concentration (4.27 ± 0.73 μM, p <1 × 10^-7^). There were no significant changes in average GA concentration in either group over the treatment period. Notably, in serum there were no significant differences between treatment groups for either MPAG, MPA or GA on day 16 (Fig. 6B). AcMPAG was not detectable in either group by our assay.

**Fig. 6.**
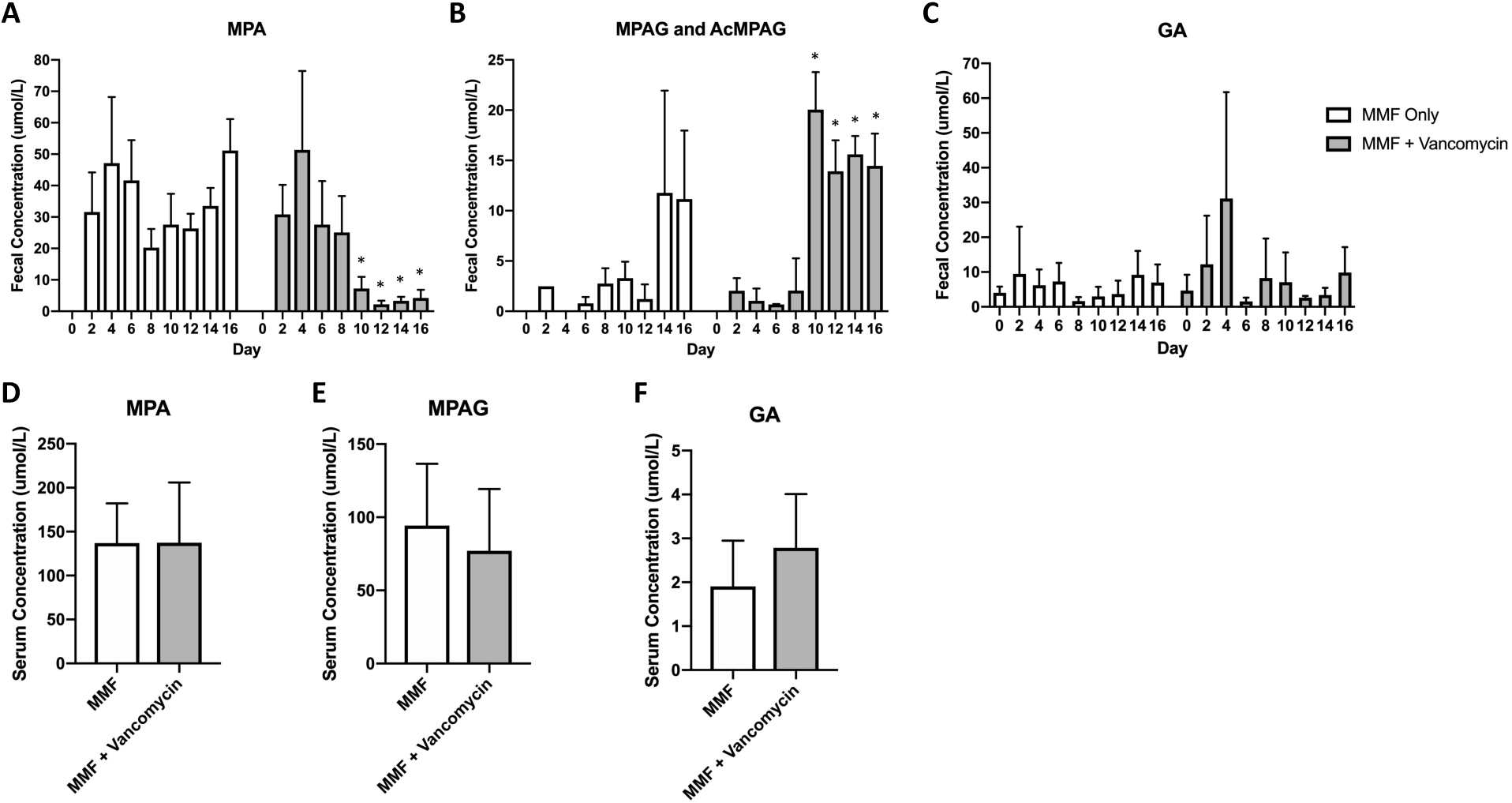
Changes in MMF metabolites. The concentration of MMF metabolites was measured in mouse fecal pellets and serum. (**A-C**) Comparison of MPA, MPAG and GA levels in mice exposed to MMF only for up to 16 days and mice initially exposed to MMF only for 8 days then with the addition of vancomycin for 8 days. Vancomycin treatment resulted in significant increases in MPAG and significant decreases in MPA levels (p < 0.05). (**D-F**) In serum from mice consuming MMF, levels of MPA, MPAG and GA were not affected after 8 days of treatment with vancomycin. AcMPAG was not detected in the serum. Data are mean ± SEM. The mean concentration of each metabolite before (days 0-8) and after (days 10-16) the introduction of vancomycin was compared independently in each treatment group using a two-way ANOVA with *p < 0.0005.

### Intrarectal administration of MPA promotes weight loss and inflammation

To elucidate the identity of the toxic metabolite, we directly administered MPA and GA into the mouse distal colon. There was a concentration-dependent effect of MPA which caused significant weight loss (p = 0.002) in treated animals and appeared to promote inflammation as indicated by a trend towards increased activity of colonic myeloperoxidase activity (Fig. S6).

### GUS activity in human stool correlates with MMF exposure

Human fecal GUS activity measured in stool samples obtained from adult and pediatric heart transplant recipients was variable but correlated with MMF exposure (Fig. 7). Patient 1, with the highest level of GUS activity, experienced significant neutropenia that was attributed to MMF. Patient 3 is the patient from our case report who had developed hematochezia and lymphonodular hyperplasia related to MMF (*19*). Patients 10 and 11 were pediatric patients whose MMF had been discontinued following the recent diagnosis of post-transplant lymphoproliferative disease.

**Fig. 7.**
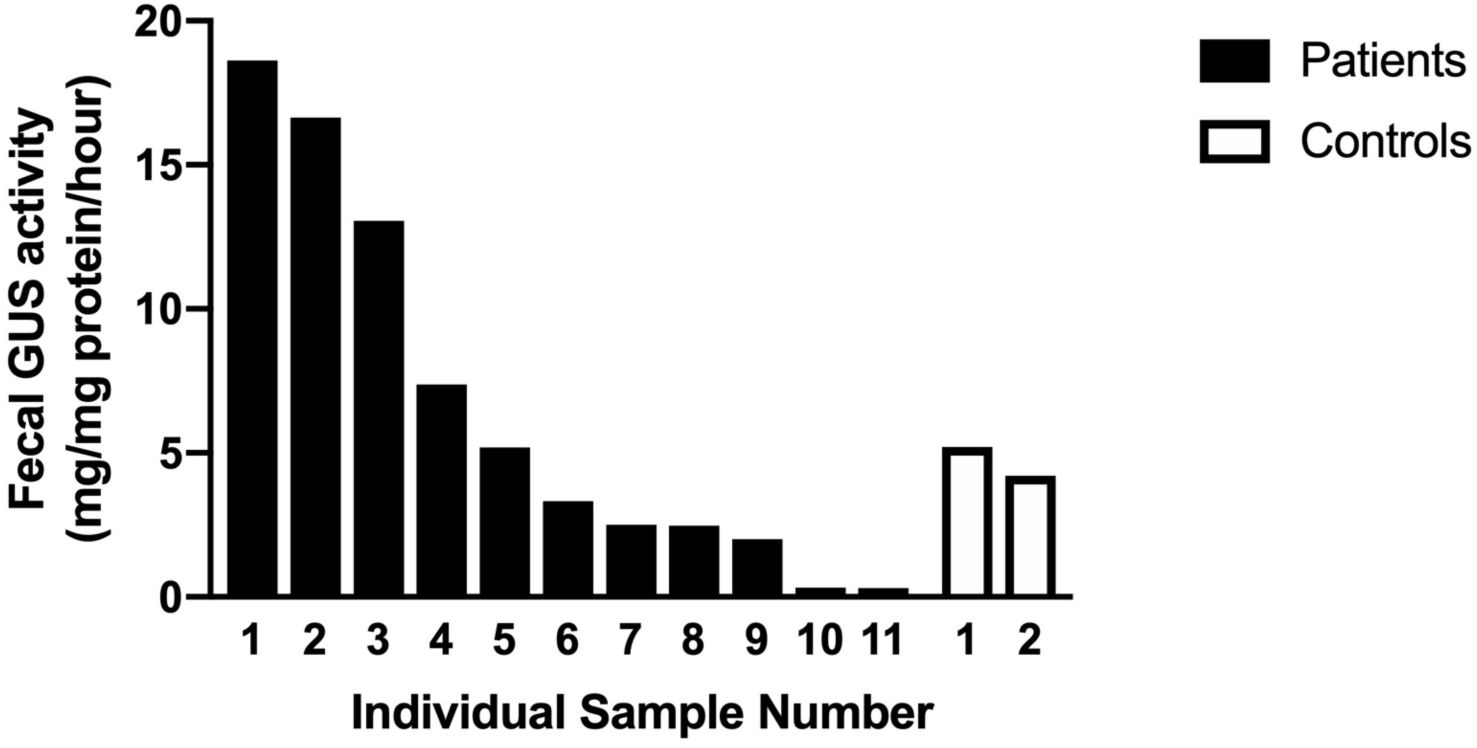
GUS activity in human stool. Samples were collected from 11 adult and pediatric heart transplant recipients and two healthy, non-transplant controls. The highest level of GUS activity seen in patient 1 correlated with lymphopenia requiring a dose reduction of her MMF. Patient 3 had developed hematochezia and colonic lymphonodular hyperplasia while taking MMF. Patients 10 and 11 had been taken off MMF prior to sample collection.

## Discussion

The use of MMF as an immunosuppressant is frequently limited by GI side effects. Attempts to relieve toxicity by modifying drug delivery, either through use of an enteric-coating or intravenous administration, have not been successful (*21-23*). The metabolism of MMF and MPA have been well described, but the mechanism of toxicity has remained elusive. Previously, we have demonstrated the requirement for an intact gut microbiome for MMF to cause GI toxicity in the mouse (*20*). Here, we now demonstrate the ability of a single antibiotic, vancomycin, to both prevent and reverse MMF-induced GI toxicity through the reduction of GUS expression, activity and GUS-producing bacteria and, consequently, decreased catabolism of the MMF metabolite MPAG to free MPA. In our mouse model, reduced fecal concentrations of MPA were associated with reduced GI toxicity.

Complete eradication of the gut microbiome with a cocktail of broad-spectrum antibiotics was able to prevent and reverse MMF-induced GI toxicity (*20*). From those four antibiotics, it appears that only vancomycin is necessary and sufficient for the prevention and reversal of MMF-induced weight loss. Vancomycin was also to reverse MMF-induced colonic inflammation as demonstrated by improvements in cecum, liver and spleen weights and colon length. Colonic tissue homogenates also showed changes in multiple cytokines and chemokines consistent with a reduction in the pro-inflammatory state induced by MMF exposure. The inflammatory mediators TNFα, IL-6, KC (CXCL1), MIP-2 (CXCL2), LIF, G-CSF and M-CSF play a role in the stimulation of acute phase proteins, granulocyte and macrophage production and the promotion of immune cell migration to the site of inflammation in response to bacterial antigens such as lipopolysaccharide (*24-26*). Reduced expression suggests that vancomycin is removing an inflammatory stimulus from the colonic lumen. This inflammatory stimulus was previously postulated to be a pathogenic bacterium (i.e. Escherichia/Shigella) that expanded due to MMF-induced dysregulation of the microbiome (*20, 27*). However, our present data suggests that the factor initiating the inflammatory response may be the free MPA, released after bacterial GUS metabolism in the lumen, which interacts with the epithelium in the distal GI tract. Evaluating how the composition and function of the gut microbial community was altered by vancomycin provided insight into this mechanism of action.

Using 16S rRNA amplicon sequencing, we determined which bacteria thrived during MMF exposure, presumably due to their consumption of free GA liberated from the catabolism of MPAG as a nutrient source (*16, 28*) and were subsequently eliminated by the addition of vancomycin. After 8 days of MMF-only treatment, there was a consistent abundance of the bacterial classes Betaproteobacteria, Bacteroidia, Bacilli, Gammaproteobacteria, Erysipelotrichia and Alphaproteobacteria. After initiating vancomycin treatment, the classes Bacteroidia, Clostridia and Erysipelotrichia were severely depleted, allowing for expansion of the classes Gammaproteobacteria and Bacilli, a response that is also observed in humans (*29*). The potential pathogenic bacteria Escherichia/Shigella (which cannot be distinguished based on 16S rRNA sequencing) of class Gammaproteobacteria, previously thought to be a primary cause of inflammation following MMF-induce dysbiosis of the microbiota (*20, 27*), expanded during vancomycin treatment while inflammation and toxicity improved, suggesting that members of the Gammaproteobacteria are not the primary driver of MMF toxicity. The gut bacteria *Bacteroides spp*. and *Roseburia spp.*, both enriched in the presence of MMF and eliminated by vancomycin, encode for GUS enzymes and have demonstrated GUS activity *in vitro* (*30-33*). It was expected that MPAG would apply a positive selective pressure to bacteria encoding MPAG-compatible GUS enzymes since the resultant GA can be used as a carbon source during anaerobic metabolism (*34*). Indeed, *Bacteroides fragilis*, *B. caccae* and *B. vulgatus* all increased in abundance during MMF treatment, suggesting they are capable of metabolizing MPAG and utilizing the resulting GA. Furthermore, these bacterial species were effectively eliminated by vancomycin, suggesting the capacity for bacterial GUS activity was reduced. This hypothesis was supported by our PICRUSt analysis and *in vivo* imaging of GUS activity.

PICRUSt predicts the functional content of a metagenome from 16S rRNA amplicon sequencing data (*35*). Functional orthologs are annotated in the Kyoto Encyclopedia of Genes and Genomes Orthology database (*36-38*). In bacterial DNA extracted from mouse fecal pellets, PICRUSt predicted an increase in the abundance of GUS orthologs in response to MMF after 4 days. This suggests a rapid increase in bacterial metabolism in response to the introduction of MMF, but this increase does not appear to be sustained. This increase in predicted gene expression was supported by our *in vitro* assay for GUS activity which showed a similar, transient increase in activity. In humans, levels of GUS activity were quite variable but did correlate with MMF-related side effects for two of the patients with the highest level of GUS activity and with the lowest levels seen in patients who had been recently taken off MMF as part of a reduction in their immunosuppression related to the diagnosis of malignancy. Levels of stool GUS activity in humans on MMF may therefore represent a potential biomarker to assess for drug-related toxicity although this remains to be tested in a larger cohort.

Assessment of bacterial GUS activity *in vivo* based upon activation of a fluorescent substrate after GUS hydrolysis (*39*), demonstrated an increase in overall bacterial GUS activity in the proximal colon, but not the cecum, of MMF-treated mice as compared to controls that was abolished by treatment with vancomycin. Evaluation of bacterial populations in the proximal colon specifically may confirm the identity of the bacteria that are responsible for MPAG catabolism and the liberation of free MPA.

We assessed the effect of a vancomycin-induced reduction in bacterial GUS activity on the hydrolysis of MPAG to MPA and GA in the GI tract of MMF-treated mice using mass spectrometry. In fecal pellets, relatively higher levels of MPA and lower levels of MPAG were measured during MMF treatment. The introduction of vancomycin induced a sharp increase in MPAG concentration, in tandem with a drastic reduction in the concentration of MPA, demonstrating that vancomycin does indeed alter bacterial metabolism of MPAG in the colon. Reduced colonic concentrations of MPA in our model coincided with reduced GI and systemic toxicity, suggesting that MPA is the major contributor to MMF-induced GI inflammation and toxicity. Importantly, the serum concentration of MPA was not significantly reduced in the vancomycin-treated animals, suggesting that MMF toxicity is occurring primarily in the GI tract, and the systemic toxicity (i.e. weight loss) is not a result of toxic doses of MPA in the blood.

Multiple human clinical reports have indirectly implicated GUS involvement in MMF metabolism (*40-42*). The induction of weight loss and colonic inflammation by the intrarectal administration of MPA and recent work by others further supports our hypothesis that MPA is responsible for the GI toxicity experienced by patients. Heischmann et al. found that MPA depleted the intracellular guanosine pool in human intestinal LS180 cells, affecting a number of cellular processes including protein, lipid and fatty acid metabolism and potentially leading to reduced membrane integrity (*43*). This effect occurred despite high levels of guanosine supplementation. Given that intestinal epithelial cells import guanosine from the GI lumen (*44*), their data suggest that MPA inhibits this process. Qasim et al. demonstrated that MPA directly modulated intestinal epithelial tight junctions in Caco-2 cell monolayers by upregulating myosin light-chain kinase leading to increased phosphorylation of myosin II regulatory light-chain and increased tight junction permeability (*45, 46*). This may be occurring via MPA-stimulated activation of the midkine-dependent PI3K pathway (*47*). Our previous work supports these observations of MPA-induced epithelial barrier dysfunction as we found that MMF exposure led to an increase in serum lipopolysaccharide, an indirect measure of intestinal epithelial barrier function (*20, 48*). Together, this data suggests that MPA may trigger local and systemic toxicity by disrupting GI epithelial barrier function, potentially allowing toxic metabolites to cross into the tissue and circulation.

While our data strongly support the expansion of GUS-expressing bacteria and alterations in the gut metabolism of MPA/MPAG in the mechanism of MMF-induced GI toxicity, we have yet to directly characterize the mechanism(s) through which this occurs. However, this work describes a clear path to further test our hypothesized mechanism. Additionally, our *in vivo* experiments represent a short-term treatment regimen of MMF, whereas exposure in the human is chronic. This difference in duration of exposure, and the observation in mice that the increase in GUS gene expression and enzyme activity appears to be relatively transient, suggest that serial samples from humans after the introduction of MMF will be very useful in evaluating our proposed mechanism of MMF toxicity. Lastly, while our data suggest that vancomycin can attenuate the MMF toxicity by targeting GUS-expressing bacteria, the non-specific devastation of the gut microbiome suggests that an alternative approach using specific GUS inhibitors may be a more effective therapeutic modality (*49*).

In summary, our data suggest that bacterial GUS metabolism of MPAG to free MPA in the colonic lumen is responsible for the GI toxicity associated with MMF use. The GUS pathway responsible for MMF metabolism affects multiple other commonly prescribed drugs including non-steroidal anti-inflammatory drugs, acetaminophen, irinotecan and cannabinoids (*50*). The ability of vancomycin to reduce toxicity suggests a possible therapeutic approach to treat or prevent MMF-induced GI toxicity.

## Materials and Methods

### Animals

All animal procedures were approved by the University of Calgary Animal Care Committee (protocol AC14-0068) and were in accordance with the Canadian Council on Animal Care policies. Female C57BL/6 mice aged 7-10 weeks (Charles River or Jackson Laboratory) were used in all experiments. Mice were housed in sterilized, filter-top cages (Techniplast) under specific pathogen-free (SPF) conditions, separated by treatment group and fed control chow (Teklad Global 16% Protein Rodent Diet TD.00217), for a minimum of 1 week before commencing any experiment. MMF-containing chow was administered *ad libitum* at a concentration of 0.563% w/w (*51*). Vancomycin (500 mg/L) was administered in the drinking water *ad libitum*. Consumption of both chow and water was monitored daily.

### Sample collection and assays

Mice were individually placed in clean, ethanol-sterilized cages and immediately upon defecation each fecal pellet was placed in a sterile collection tube, snap-frozen and stored at - 80°C. Whole blood was collected from anesthetized animals by cardiac puncture. Serum was isolated by centrifugation at 1500*g* × 15 mins and stored at −80°C. Liver, spleen and intestine were collected immediately after euthanasia. Each tissue was weighed, and colon length was measured before being placed in microcentrifuge tubes, snap-frozen and then stored at −80°C. Hematocrit was determined using a micro-hematocrit capillary tube reader. Whole colonic tissue was homogenized in cell lysis buffer and protein concentration was determined using the Precision Red Protein Assay (Cytoskeleton). Samples were sent to Eve Technologies for analysis using their 32-multiplex Mouse Cytokine/Chemokine Array.

### Bacterial community analysis

Bacterial genomic DNA was isolated from mouse fecal pellets (*52, 53*) and concentration determined using a Qubit 2.0 fluorometer (Invitrogen). Variable regions 3 and 4 of the 16S rRNA gene were amplified by PCR using barcoded primers (*54*) before sequencing on an Illumina MiSeq in the Nicole Perkins Microbial Communities Core Laboratory at The University of Calgary. Primers and poor-quality reads were removed using Cutadapt v1.16 (*55*) then processed using DADA2 v1.6.0 (*56, 57*). Taxonomy was assigned using the Silva Taxonomy Training Set v128 and the Silva Species Assignment Set v128 (*58*). Visual and statistical analysis of amplicon sequencing data was performed using the R packages Phyloseq (*59*) and Vegan v2.5-2 with support from dplyr 0.7.5 and ggplot2 v3.0.0. Reads of non-bacterial origin and samples with insufficient depth (<10,000 reads) were removed. Observed species, Shannon diversity index, and Simpson’s index were determined. Multiple indices were used to ensure a potential bias did not arise from unequal numbers of reads between samples (*60*). Beta-diversity was assessed using a non-metric multidimensional scaling (NMDS) plot of Bray-Curtis dissimilarity. The R package DESeq2 v1.18.1 and microbiomeSeq v0.1 were used to calculate significantly differentially abundant taxa in the vancomycin-treated group over time. Prior to further analysis and visualization of taxa specific differences between samples or treatment groups, reads in each sample were normalized to relative abundance.

For input into PICRUSt (Phylogenetic Investigation of Communities using Reconstruction of Unobserved States) v1.1.3 (*35*), reads were reprocessed into operational taxonomic units (OTUs) (*56*). Read quality was assessed using FastQC v0.11.5 and paired-end reads were merged and filtered using PEAR v0.9.10 (*61*). Chimeric sequences were removed using VSEARCH v1.11 (*62*). Sequences were clustered into OTUs using a closed reference OTU picking pipeline using QIIME v1.91 (*63*) and mapped to the Greengenes Database v13_8 (*64*). Within PICRUSt, the OTU table was 16S rRNA copy number normalized and then functional predictions of KEGG Orthologs (KOs) were predicted using PICRUSt’s Hidden State Prediction (HSP) algorithm.

### Ex vivo imaging of GUS activity

We performed *ex vivo* tissue imaging of intestinal bacterial GUS activity through quantification of fluorescence following hydrolysis of fluorescein di-b-D-glucuronide (FDGlcU, Sigma-Aldrich) to fluorescein (*65*). Three hours prior to imaging, each animal was gavaged with FDGlcU (7.3 μmol/kg). Immediately before imaging, mice were euthanized and abdominal organs were excised and imaged using an In Vivo Xtreme 4MP imaging platform (Bruker). Reflectance imaging (2 second exposure time) was followed by fluorescent imaging (5 second exposure time, 470 nm excitation and 535 nm emission filters) and X-ray imaging (10 second exposure time). Pixel binning was kept constant at 4×4. Images were acquired and analyzed using the Bruker molecular imaging software MI SE v7.1.3.20550. GUS activity was quantified by measuring the mean fluorescence after background subtraction in a given region of interest which was kept constant over the time period of imaging.

### Metabolomics

Metabolites were extracted from fecal pellets using ice-cold 50% methanol followed by homogenization with ceramic beads using a BeadRuptor 4 (Omni) and centrifugation at 10,000*g* × 15 mins at 4°C. Following removal of the supernatant, the process was repeated and the supernatants pooled. Whole blood was mixed with 50% MeOH and centrifuged at 14,600*g* × 5 mins. Metabolites in the supernatant were measured using ultra-high performance liquid chromatography (UHPLC) on a Vanquish UHPLC platform (ThermoFisher Scientific) using either hydrophilic interaction liquid chromatography (HILIC) and a Syncronis HILIC LC column (ThermoFisher Scientific) for fecal metabolites or a reverse phase octadecyl carbon chain bonded silica (C18) and an Accucore Vanquish C18+ UHPLC column (ThermoFisher Scientific) for serum metabolites. Mass spectrometry (MS) was conducted using a Q Exactive HF Hybrid Quadrupole-Orbitrap Mass Spectrometer (ThermoFisher Scientific) in negative ion full scan mode. Raw data acquisition was carried out using Xcalibur 4.0.27.19 software. Peak annotation was performed using MAVEN (*66, 67*) and ElMaven (*68*). Each targeted metabolite was aligned with a corresponding standard and standard curve, run concurrently with the samples, to validate identification and determine concentration.

### Enzymatic assay for GUS activity

Fecal pellets were homogenized in PBS (0.01 M, pH 7.0) using a BeadRuptor 4 (Omni) then sonicated using a Branson 3510 Water-Bath Ultrasonicator. Activity of extracted bacterial GUS was measured by hydrolysis of phenolphthalein glucuronide (Sigma-Aldrich) to free phenolphthalein and quantified by spectrophotometry (*69*). GUS activity was calculated as mg of free phenolphthalein liberated/mg protein/hour of incubation.

### Intrarectal administration of MPA and GA

Injections were performed daily for 8 days using a 1 mL syringe and a polyethylene catheter (PE-50). With the mouse immobilized, the catheter was inserted 3-4 cm intrarectally and 200 μL of either vehicle (50% DMSO), GA (17.5 mM) or MPA (1.75 mM or 17.5 mM) was slowly injected by hand. After removal of the catheter, the rectum was held closed for a minimum of 2 minutes. Body weight of each animal was recorded daily. After 8 days, animals were sacrificed. Colonic tissue was homogenized in hexadecyltrimethylammonium buffer (HTAB) using a Polytron homogenizer (Kinematica). O-Dianisidine (Sigma-Aldrich) was added to the cell lysate and its oxidation was measured by absorbance at 450 nm using a SpectraMax Plus 384 microplate reader. Myeloperoxidase (MPO) activity (units/mg tissue) was calculated from absorbance based on a standard curve.

### Statistical analysis

All statistical analysis was conducted using R and Prism 7 (GraphPad). Data are expressed as mean ± standard deviation. For each set of data, normality was determined using the Shapiro-Wilk Normality Test. For data with 2 treatment groups, each experimental group mean was compared pairwise against the control group mean using either Welch’s t-test or Wilcoxon Rank Sum test with a correction for multiple comparisons using the Benjamini-Hochberg procedure. Means of parametric data with >2 treatment variables compared at a single time point were analyzed using a one-way ANOVA. Significant ANOVA results were followed by the Tukey Honest Significant Differences post hoc test. Means of non-parametric data with >2 treatment variables compared at a single time were analyzed using pairwise Wilcoxon Rank Sum Tests with p-value correction for multiple testing using the Benjamini-Hochberg procedure. To compare means of two treatment groups over time, a two-way ANOVA was used, followed by the Tukey Honest Significant Differences post hoc test. If the two-way ANOVA residuals were non-parametric, the Kruskal-Wallis Rank Sum Test followed by Dunn’s Kruskal-Wallis Multiple

Comparisons post hoc test was used. Intergroup significance in β-diversity was computed using the Adonis test applied pairwise, using the Benjamini-Hochberg procedure for p-value correction. A p value <0.05 was considered significant except for the assessment of colonic tissue cytokines and chemokines, the pairwise group comparisons of observed species for α- diversity and β-diversity comparisons between MMF- and vancomycin-treated animals, where a p value <0.1 was considered significant.

## Acknowledgments

The authors acknowledge the assistance of Dr. Bjoern Petri and the Mouse Phenomics Resource Laboratory in the Snyder Institute for Chronic Diseases at the University of Calgary with the *in vivo* mouse imaging experiments.

## Author contributions

The study was designed by MRT, KLF, SAH and SCG. Experiments were performed by MRT, KLF, HR and AM. Data was analyzed by MRT and SCG. MRT and IAL performed the MS assays and analysis. MRT and SCG wrote the manuscript. All authors reviewed and approved the manuscript.

## supplementary-materials

**Fig. S1.**
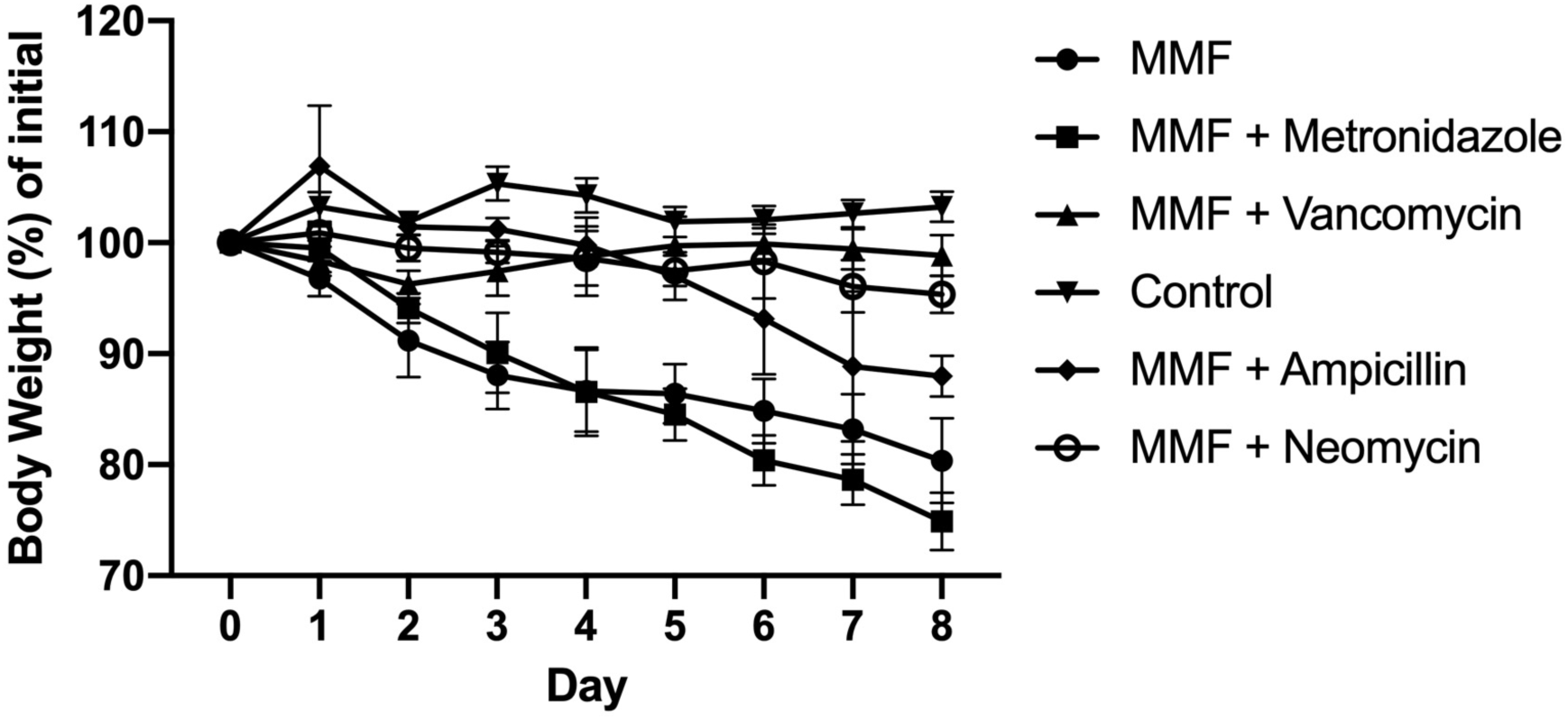
Effect of individual antibiotics (ampicillin, neomycin, metronidazole or vancomycin) on mouse body weight in the presence of MMF. Control animals received normal chow and water. Data are shown as mean and standard deviation for each day.

**Fig. S2.**
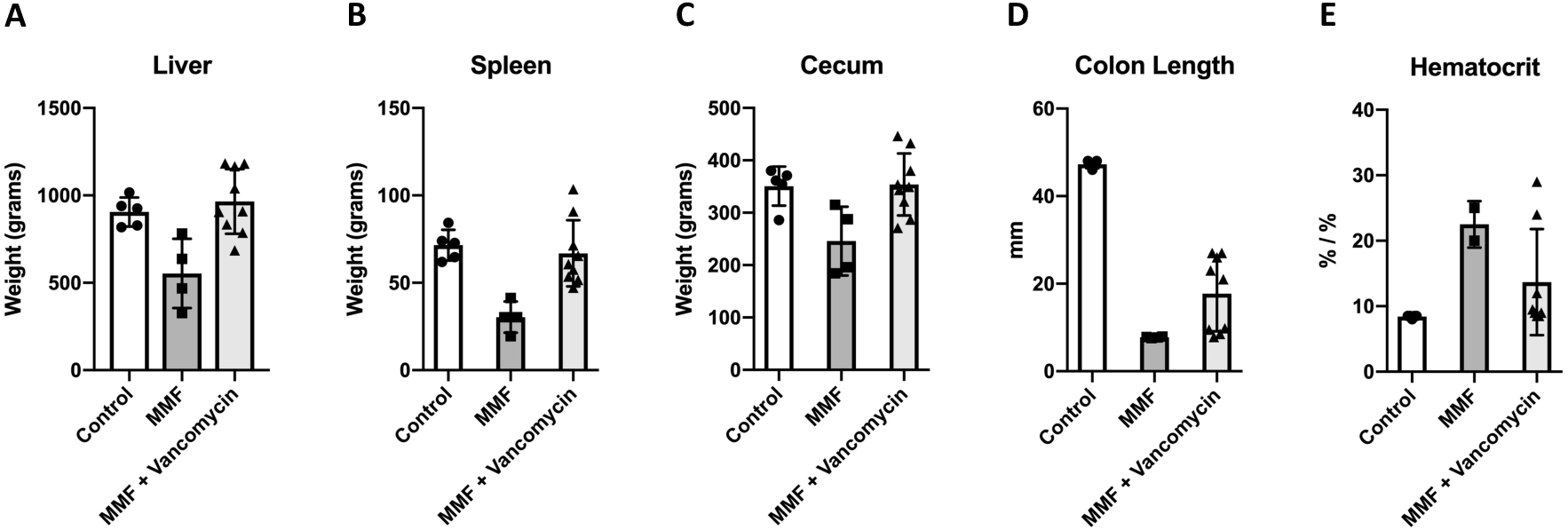
Effect of 8 days of exposure to MMF alone or MMF with vancomycin on weight of (**A**) liver, (**B**) cecum, liver and (**C**) spleen. Effect of MMF and MMF with vancomycin after 8 days on (**D**) colon length and (**E**) hematocrit. Control animals received unmedicated chow and plain water. Data are shown as mean and standard deviation.

**Fig. S3.**
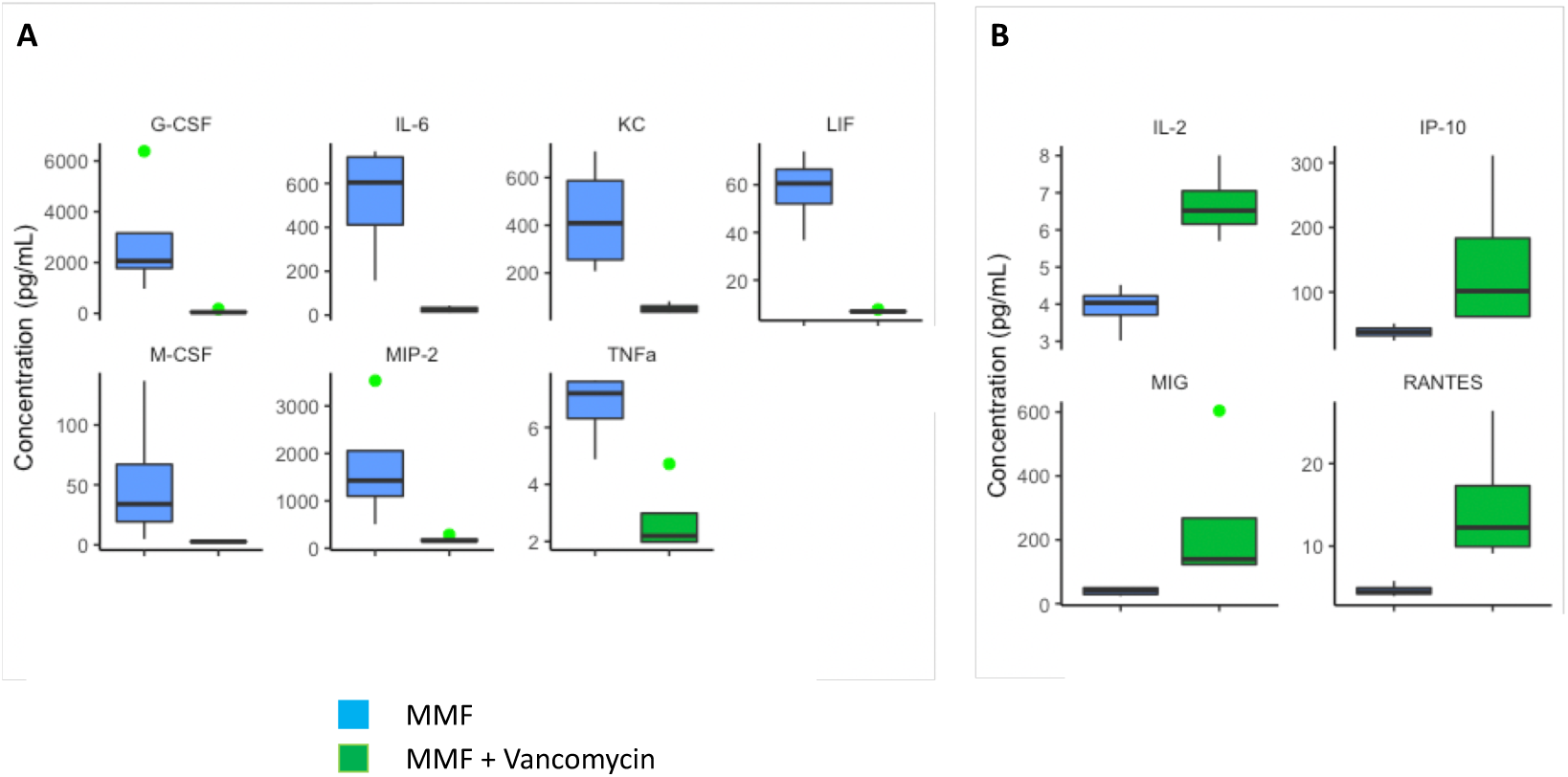
Effect of 8 days of exposure to MMF alone or MMF with vancomycin on individual cytokines and chemokines. (**A**) Vancomycin induced a significant decrease in seven cytokines. (**B**) Vancomycin resulted in a significant increase in four cytokines.

**Fig. S4.**
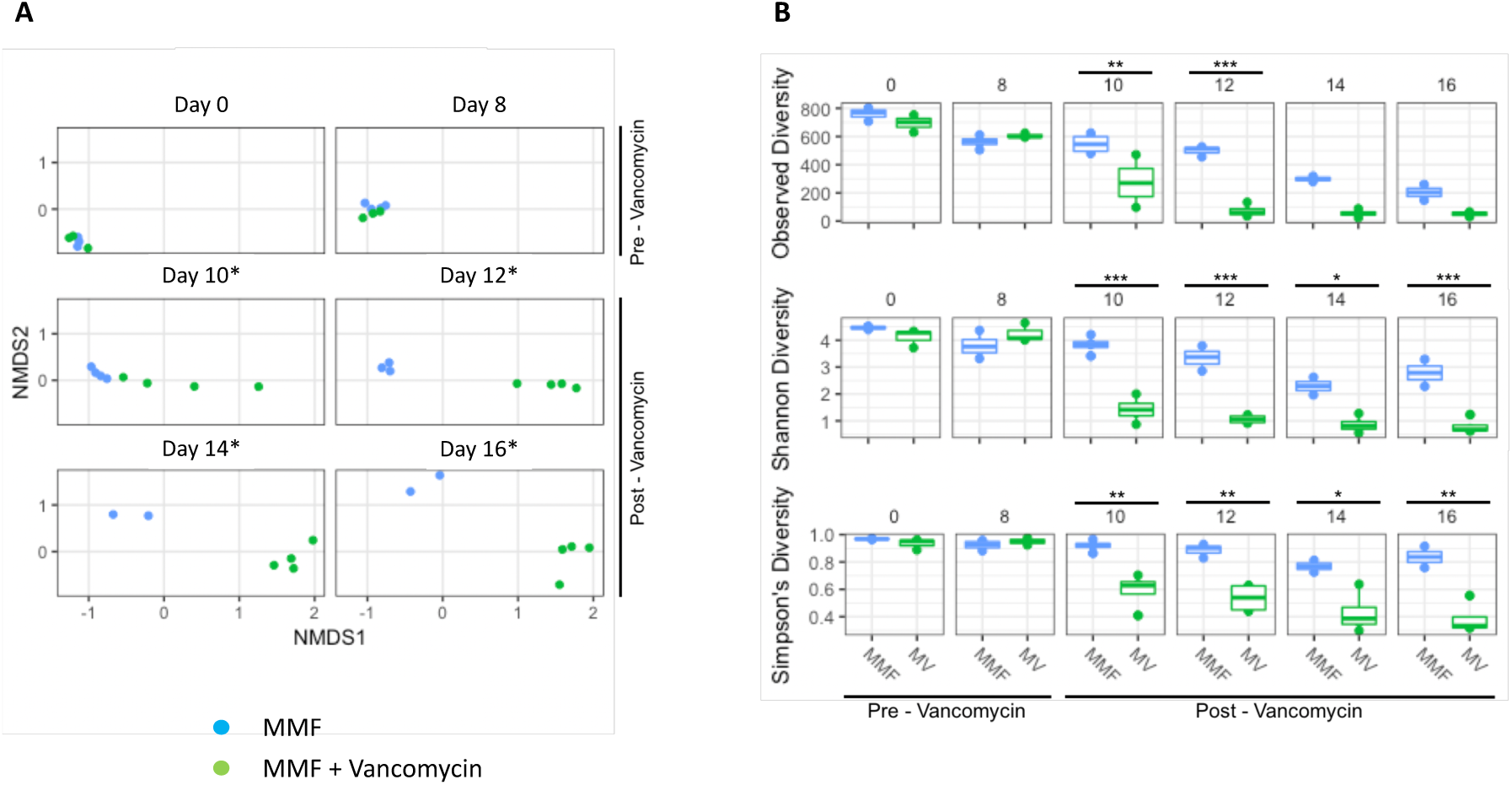
(**A**) Non-metric multidimensional scaling (NMDS) plots for β-diversity (Bray-Curtis dissimilarity) for bacteria in fecal pellets collected from mice exposed to MMF only (n=2-4) or MMF with vancomycin (n=4 individual animals). Community composition was determined using 16S rRNA amplicon sequencing of bacterial DNA extracted from fecal pellets. Each data point represents an individual sample. Intergroup comparisons were performed for each time point using Adonis with *p <0.1. (**B**) Effect of MMF alone (MMF) and MMF with vancomycin (MV) exposure on bacterial α-diversity for three different indices (Observed, Shannon and Simpson’s). Community composition was determined using 16S rRNA amplicon sequencing from mouse fecal pellets collected from 2-4 individual animals. Significant differences in diversity between groups was determined using a two-way ANOVA with *p <0.05, **p < 0.001, ***p < 0.0005.

**Fig. S5.**
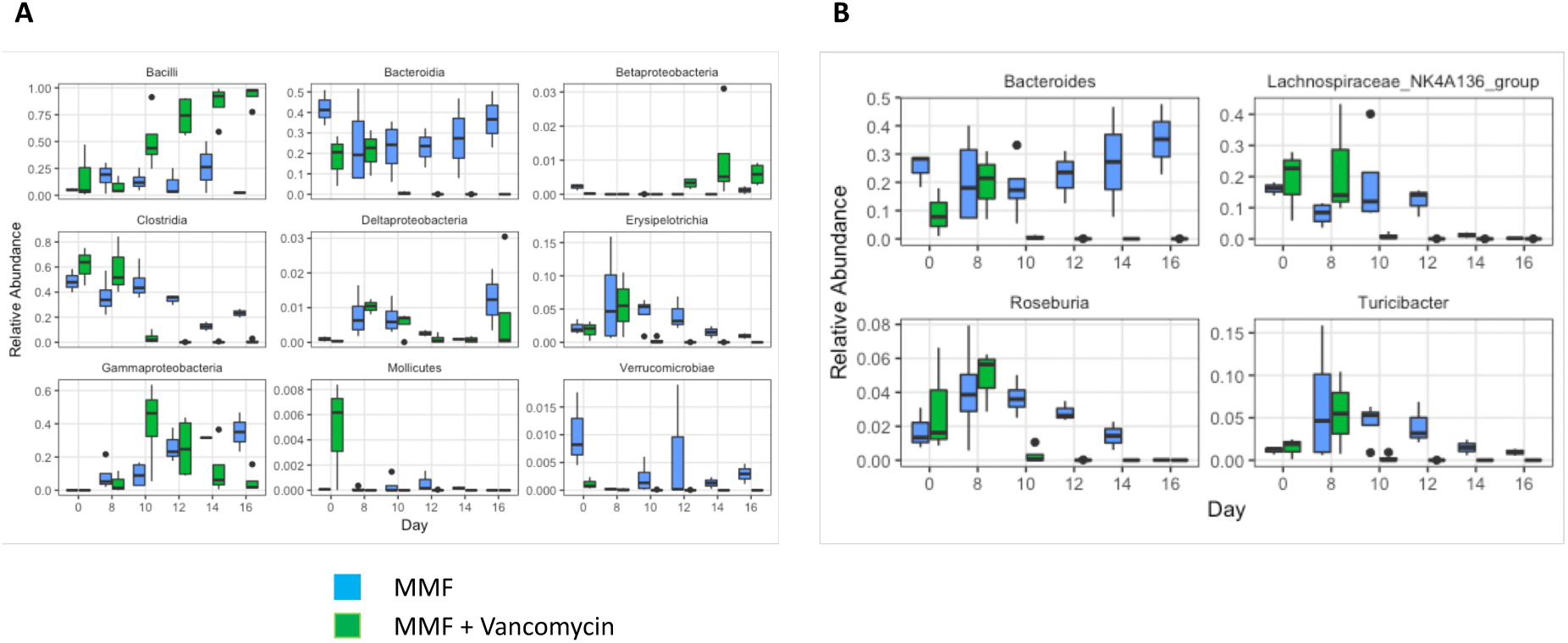
(**A**) Effect of MMF and vancomycin (introduced on day 8) on relative abundances of bacterial classes in mouse fecal pellets. (**B**) Changes in relative abundance of bacterial genera that showed differential abundances in the absence and presence of vancomycin.

**Fig. S6.**
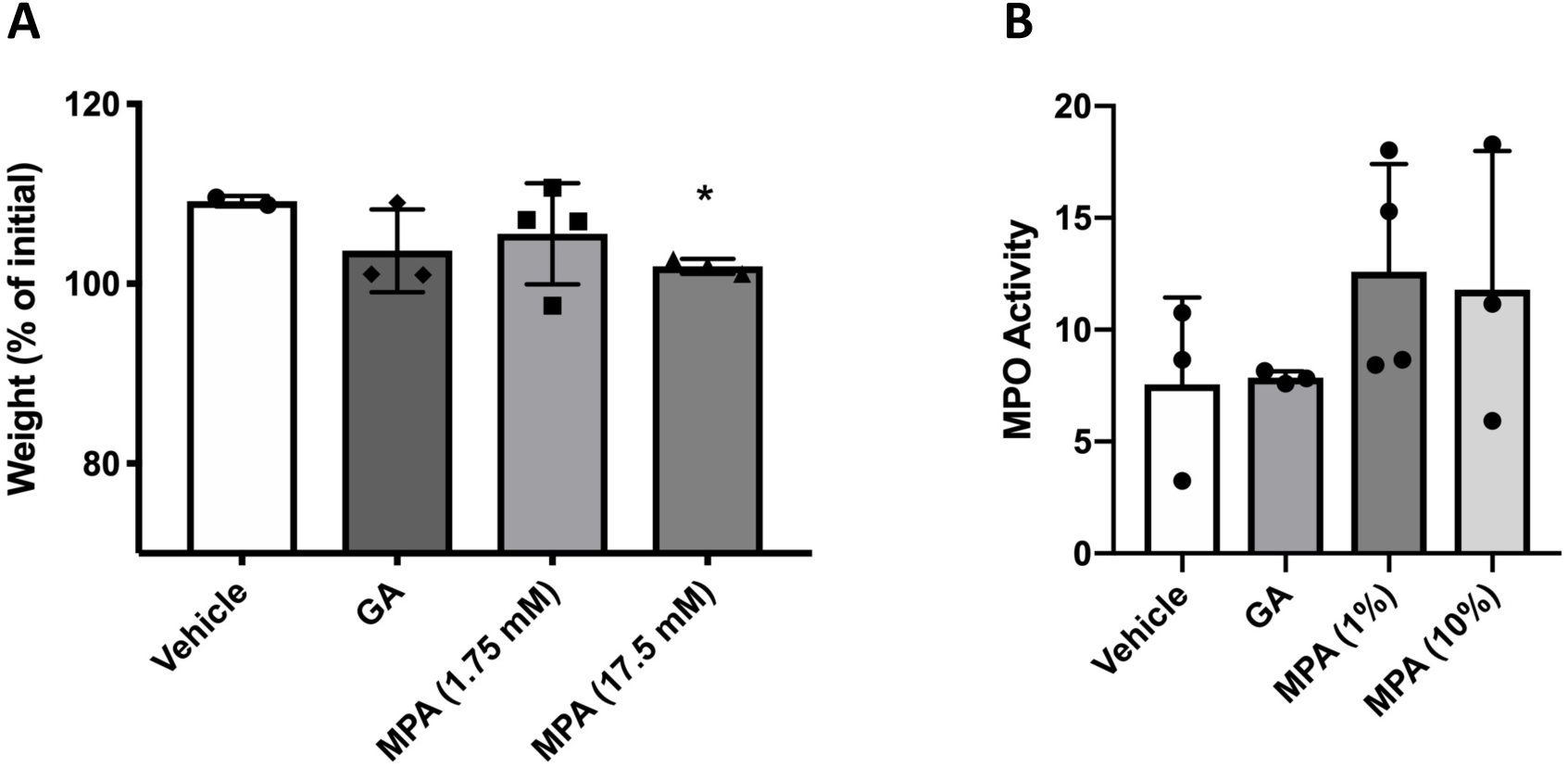
(**A**) Effect of daily intrarectal administration of MPA (1.75 mM and 17.5 mM concentrations) and GA (17.5 mM) on mouse body weight after 8 days. Only 17.5 mM MPA exposure resulted in significant weight loss (*p = 0.002). (**B**) Effect of daily intrarectal administration of MPA (1% and 10% concentrations) and GA on MPO activity in whole colonic homogenates. There was a positive trend towards increased MPO activity (units/mg tissue) in mice treated with MPA but not GA. Data are shown as mean and standard deviation.

